# Integrating supervised and unsupervised machine learning for behavior segmentation reveals latent frailty signatures and improves aging clocks in isogenic and outbred mice

**DOI:** 10.64898/2026.03.23.713050

**Authors:** Gautam S. Sabnis, Dustin M. Miao, Vivek Kumar

## Abstract

Manual frailty index (FI) assessment in mice is a strong predictor of morbidity and mortality, and is frequently used in mechanistic and translational geroscience. However, it is labor-intensive, requires expert training, and is vulnerable to scorer variability. We previously developed a visual frailty index (vFI) that objectively predicts age and frailty using expert-defined, supervised behavioral features extracted from open-field videos. However, relying solely on human-defined features may miss subtle, latent behavioral signatures of aging. Here, we test whether unsupervised behavioral discovery using Keypoint-MoSeq (KPMS) could uncover these hidden signatures and improve the prediction of aging-related outcomes. Using a large dataset of isogenic C57BL/6J (B6J) and genetically diverse Diversity Outbred (DO) mice, we find that unsupervised features are highly predictive of chronological age, biological frailty, and the proportion of life lived. Notably, while supervised features overall outperformed unsupervised features in these tasks, combining both feature sets yielded the highest predictive accuracy across all outcomes. Despite these improvements, models trained on either feature set failed to generalize across strains, confirming that behavioral manifestations of aging are strongly population-specific. These findings demonstrate that supervised and unsupervised machine vision provide complementary information, establishing a highly sensitive, scalable, and non-invasive framework for objective and scalable geroscience in rodents.

## 2 Introduction

Aging is a universal, gradual process that affects nearly every biological system, but it occurs at different rates in different individuals^1, 2^. This heterogeneity in biological aging is captured by the concept of frailty, defined as an increased vulnerability to adverse health outcomes^3^. In preclinical research, the mouse frailty index (FI) is an invaluable tool for quantifying this biological age. During a manual FI assessment, a trained technician evaluates a mouse across a set of age-related health deficitsscoring each item as 0 (absent), 0.5 (partially present), or 1 (present)which are then summed to produce a cumulative frailty index score (CFI)^2, 4, 5, 6, 7, 8, 9^. Manual frailty index assessments are strong predictors of mortality and morbidity in mice, frequently outperforming other measures of biological age such as DNA methylation clocks^3, 10, 11, 12, 13^.

Despite its predictive value, the manual frailty assessment is labor-intensive, requires extensive expert training, and is highly susceptible to inter- and intra-rater variability^3, 7, 8, 9^. This scorer variability represents a critical bottleneck for the scalability and cross-laboratory reproducibility of aging studies. For example, in previous large-scale evaluations, tester effects accounted for a staggering 42% of the variance in manual FI scores for C57BL/6J (B6J) mice^14^ and 18% of the variance in Diversity Outbred (DO) mice^15^. Notably, this technical variation can sometimes exceed the biological variation introduced by severe lifespan-extending interventions, such as caloric restriction.

To directly address these limitations, we previously developed the visual Frailty Index (vFI)^14, 15^, an automated, machine-vision-based framework that predicts frailty from video recordings of mice in an open-field assay. We trained regression models that predict the CFI with high precision (1.08 ± 0.05 in B6J and 1.53 ± 0.18 in DO mice). Intuitively, an error of 1, compared to manual frailty assessment, is equivalent to 1 FI item being mis-scored by 1 point or 2 FI items being mis-scored by 0.5. The success of the vFI demonstrates that vital information regarding both chronological and biological age is deeply embedded within an animal’s spontaneous behavior. In constructing these initial aging clocks, we relied on supervised behavioral feature extraction. We trained models to specifically look for expert-defined, biologically grounded metrics, including spatial and temporal gait parameters^16^, postural changes (e.g., flexibility and spinal mobility)^14^, and traditional open-field activity measures^17^.

However, a fundamental drawback of this top-down, supervised approach is its anthropocentric bias: we only look for the behaviors we already know to associate with aging. Consequently, we may be missing subtle, non-canonical signatures of physical decline. Furthermore, supervised behavior classifiers only annotate a small percentage of the total video. Because the majority of the video data does not contain these pre-defined “behaviors of interest,” a vast amount of potentially informative high-dimensional data is discarded and left unanalyzed.

To overcome these limitations, the emerging field of computational neuroethology^18, 19^ has developed sophisticated methods for unsupervised behavior segmentation^20, 21, 22, 23, 24^. Data-driven algorithms, such as Keypoint-MoSeq (KPMS)^24^, an extension of MoSeq^23^, map the latent structure of pose trajectories without requiring manual labels. A major advantage of this approach is that it generates an unbiased, comprehensive ethogram by assigning every frame of a video to a discrete behavioral “syllable”. By utilizing 100% of the video data, unsupervised segmentation opens the door to discovering latent behavioral motifs that human observers might overlook^25^. Nevertheless, this unsupervised approach carries distinct disadvantages: it frequently extracts sub-second, highly abstract motifs or latent embeddings that are noisy and exceptionally difficult to interpret in a traditional biological context.

In this study, we sought to determine how well unsupervised behavioral features perform compared to our established supervised methods in predicting aging outcomes. Using a published large, combined dataset of B6J and DO mice, we found that unsupervised methods alone perform remarkably well. However, when unsupervised features are combined with our supervised feature set, we achieve significantly higher predictive performance across chronological age, biological frailty, and the proportion of life lived. Permutation-based model selection reveals that supervised features remain highly preferred by the models, acting synergistically alongside a targeted subset of unsupervised features, such as specific KPMS syllable usage. These results indicate that latent behavioral signatures extracted via unsupervised segmentation, when fused with targeted supervised behavioral metrics, yield a vastly more robust, sensitive, and comprehensive age and frailty clock. Our current work advances the field of automated frailty assessment by (1) Improving the current automated chronological and biological age predictors, (2) Providing the community unsupervised embeddings trained on genetically diverse mice, and (3) improving interpretability of unsupervised behavior based clocks for geroscience.

## 3 Results

### 3.1 Data collection and study design

We reanalyzed data from two previously published studies: C57BL/6J (B6J) mice from Hession et al. (14), and Diversity Outbred (DO) mice from Sabnis, Churchill, Kumar (15). The combined dataset included mice of both sexes and both strains, housed at the JAX animal facility (Supplementary Table S1). DO mice were maintained on five diets: *ad libitum* feeding (AL), fasting one day per week (1D), fasting two consecutive days per week (2D), and caloric restriction at 20% (20) or 40% (40) of baseline ad libitum food intake. In the original studies, a top-down video of each mouse was recorded during a 1-h open-field session following published protocols (see Section 5), and a trained expert scored and assigned a manual FI score after each recording.

Unsupervised features were obtained by training Keypoint-MoSeq (KPMS) on a subsample of the full dataset and running inference on all videos (Figure 1A). Sample sizes were largest for *ad libitum* feeding, with fewer animals in the fasting and caloric restriction arms; DO mice were represented across all five diet conditions (Figure 1C). The analytical framework comprised three input streams—manual frailty assessments, supervised video-derived features, and unsupervised video-derived features— feeding into models that predict biological age (FI), chronological age, and proportion of life lived (Figure 1B). Cumulative frailty index (CFI) increased with age in both B6J and DO populations, with piece-wise linear fits capturing the rise across the lifespan and residual variation at a given age reflecting diet and strain (Figure 1D). The range of distribution of cumulative frailty index (CFI) is largely identical between B6J and DO mice (Figure 1E). Our study design supports testing whether unsupervised features complement supervised features for predicting age and frailty across genetically distinct populations and dietary interventions.

**Figure 1:**
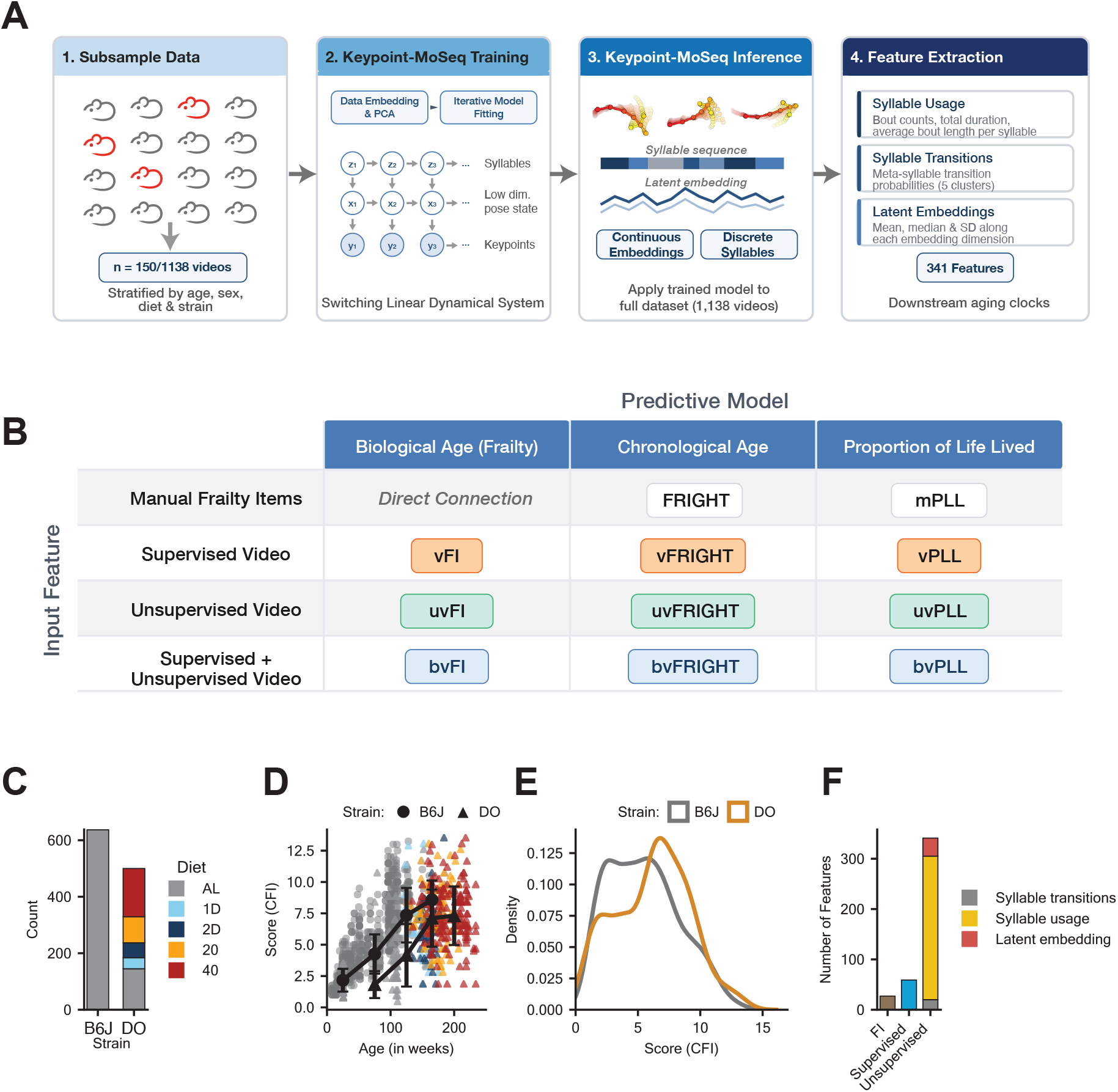
Overview of study and data distribution. (A) Diagram of KPMS processing pipeline. The full video dataset is sampled and used to train a KPMS model; the model is then used to infer on the entirety of the dataset, and features are extracted for downstream frailty, age, and PLL prediction. (B) Description of our models. We have three sets of inputs: manual frailty assessments (m), supervised video features (v), and unsupervised video features (uv). Both the supervised and unsupervised video features were extracted from an open-field assay video. In addition to the 5 models published in^26^ and^27^, we present 3 models using unsupervised video features to predict biological age (Frailty Index, FI), chronological age, and proportion of life lived (PLL). (C) Summary of the number of animals per strain and diet. The five diet intervention groups: ad libitum (AL); one day (1D) or two consecutive days (2D) per week fasting; or calorie restriction (CR) at 20% (20) or 40% (40) of estimated adult food intake. (D) Distribution of cumulative frailty index score (CFI) score by age. The black lines show piece-wise linear fits to the data separately for the two populations (n(B6J) = 126, 191, 283, 45, 0; n(DO) = 31, 57, 108, 209, 95). The center point is the mean, and the error bars are the s.d. values. Color and shape indicate diet and mouse population, respectively. (E) The cumulative frailty index score (CFI) or frailty index score (FI) distribution across B6J and DO populations. (F) The breakdown of features across the three input sets. Syllable transition represents the features derived from transition probabilities between syllables; syllable usage represents the features derived from the discrete syllable; latent embedding represents features derived from the continuous low-dimensional embedding of the animal’s motion.

### 3.2 Open-field video feature extraction and exploratory data analysis

We extracted two feature sets: supervised (S) and unsupervised (U). We previously described our open-field video data processing and supervised feature-extraction workflow. Overall, we computed 59 supervised features, categorized as open-field, gait, and morphometric-engineered features^26^. To extract unsupervised features, we trained the Keypoint-MoSeq (KPMS) model on a dataset containing 150 B6J-DO mice (Table 1). Methodological limitations of KPMS constrain the size of the training set; we therefore subsampled from the full dataset rather than training on all available videos. To ensure a balanced representation of both chronological and biological ages, and sexes in the training data, we used stratified sampling to select 150 videos (Figure S1A). Next, we ran inference with the trained model on the remaining B6J and DO mice to extract unsupervised features for all animals. The output was a syllable assignment and a continuous latent embedding for each frame of every video. The latent embedding is a low-dimensional, continuous representation of pose that incorporates time-series information across the whole video. We used these outputs to extract a total of 341 unsupervised features across the entire dataset, providing a comprehensive representation of the high-dimensional behavioral space.

**Table 1:** Sample distribution across the full dataset and the KPMS training subset for the combined model.

|  | Strain |  | Sex |  | Diet |  |  |  |  | Age (weeks) |
| --- | --- | --- | --- | --- | --- | --- | --- | --- | --- | --- |
| | B6J | DO | Male | Female | AL | 1D | 2D | 40 | 20 | Mean ± SD |
| Full dataset | 638 | 500 | 386 | 752 | 783 | 39 | 53 | 171 | 92 | 104.6 ± 61.0 |
| KPMS training | 75 | 75 | 46 | 104 | 96 | 6 | 8 | 26 | 14 | 108.7 ± 61.5 |

We derived three classes of features from KPMS inference: **syllable usage, syllable transitions**, and **latent embeddings** (Figure 1F). *Syllable usage* features were derived from the discrete syllable sequence outputted by the KPMS model. For each syllable, we extracted the number of bouts, total duration in frames, and average bout length. Due to the large number of syllables, we found that the raw syllable transition matrix was sparse and susceptible to noise (Figure S1C). To enhance the information contained within the *syllable transition* probabilities, we employed a hierarchical clustering approach to group individual syllables into five ‘meta-syllables (see Methods). These aggregated transition probabilities were then extracted to provide feature set containing syllable transitions for our downstream predictive models. Finally, we summarize the *latent embeddings* by taking the mean, median, and standard deviation along each embedding dimension. While the direct biological interpretability of these embeddings is less intuitive than that of stereotyped syllables or their transitions, we included metrics derived from the latent embeddings to evaluate whether they contain additional predictive information related to aging. The full workflow of KPMS training, inference, and feature extraction is visualized in Figure 1A.

A central objective of this study is to benchmark the performance of targeted aging-related supervised features against the behavioral repertoire extracted via KPMS. To this end, we asked whether supervised features, benefiting from expert-driven top down selection, yield fewer but higher-quality features than bottom-up discovery of KPMS. We defined “high-quality” features as those maintaining an absolute correlation coefficient greater than 0.3 (|*r*| > 0.3) with either chronological age or manual frailty scores. To evaluate this, we calculated correlations for both B6J and DO datasets separately and plotted the empirical distributions of the correlations by feature type (Figure 2A, B). The high-quality features are indicated by rugs falling outside the |*r*| = 0.3 threshold. We found that unsupervised syllable usage features contributed the most high-quality features, especially in the DO dataset (Figure 2A, right panel). In the B6J cohort, the contributions from supervised and syllable usage features were qualitatively comparable (Figure 2A, left panel). For frailty, the highest-quality contributions shifted between feature sets: B6J showed strong signals from supervised and latent embedding features, while DO was dominated by supervised and syllable usage features (Figure 2B). Overall, these results suggest that both supervised and unsupervised features capture a larger number of high-quality signatures of aging and frailty. However, the unsupervised features, as observed with supervised features in previous work, showed a high degree of multicollinearity (Supplementary Figure S1B), suggesting that the amount of independent information captured by the unsupervised feature sets may be lower than the raw count of high-quality correlations implies.

**Figure 2:**
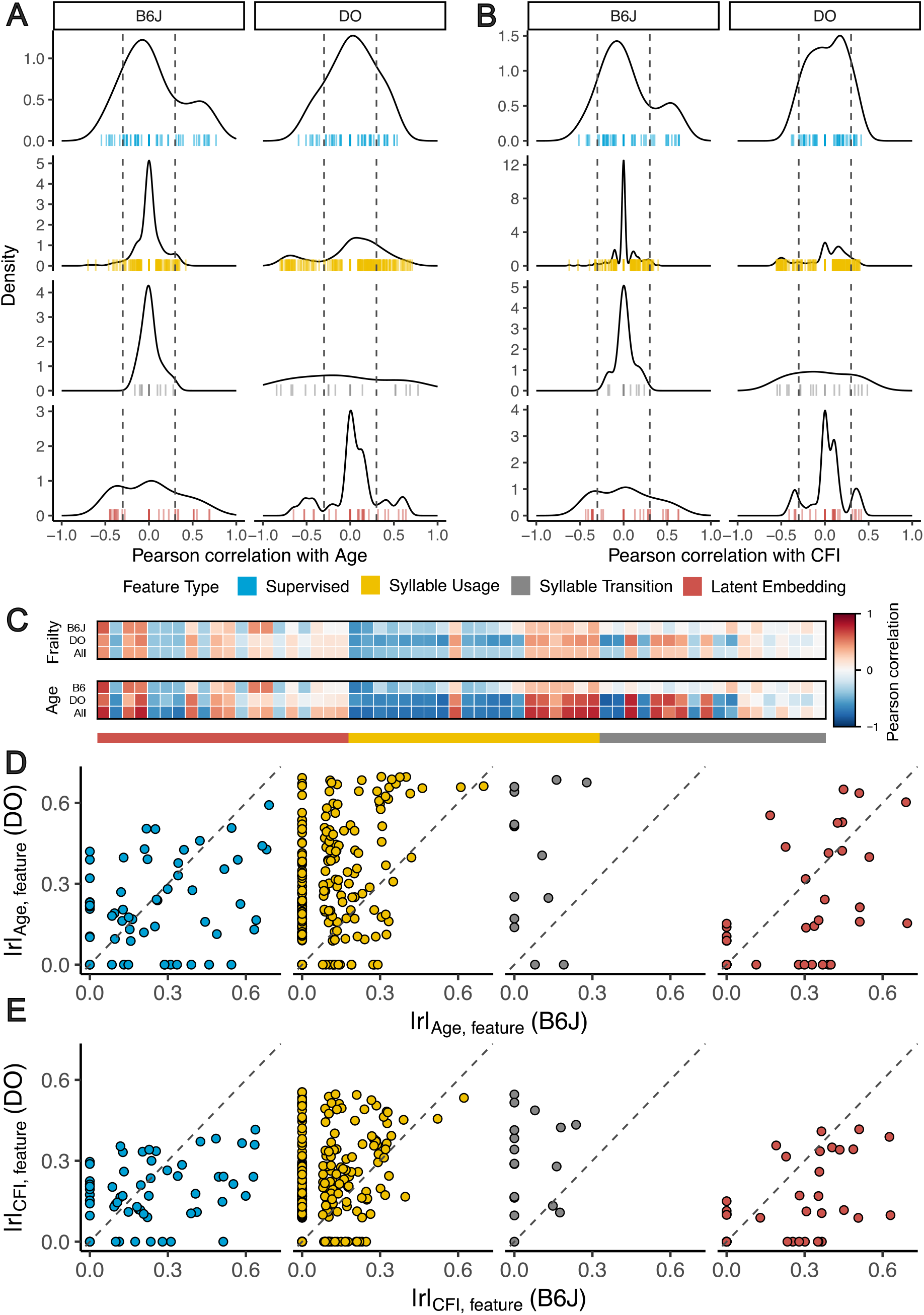
Correlation-based analysis of unsupervised video features. (A,B) Density plot of each feature category’s correlation to age (A) and frailty (B) for each strain on the combined dataset. Each colored line represents an individual feature correlation. The vertical dashed lines represent the “highly correlating” threshold at ±0.3. (C) Heatmap of Pearson correlations of the top 20 highest-correlating unsupervised features to frailty score (CFI) to CFI (top) and chronological age (bottom). (D,E) Scatterplots showing strain-specific correlation to age (D) and frailty (E) for each feature category for B6J (*x*-axis) and DO (*y*-axis). Points that lie on the dashed diagonal lines represent features that correlate equally well with both

We analyzed correlations among all features within each feature set, assessed separately for age and frailty (Figure 2C). We observed both conserved and distinct correlation patterns across the B6J, DO, and B6J-DO (all) datasets. To investigate these patterns further, we constructed pairwise scatterplots comparing B6J-DO correlations for both supervised and unsupervised feature sets (Figure 2D,E). We found that a significantly larger proportion of features derived from syllable assignments (both usage and transitions) were DO-specific (aligned along the *y*-axis) than B6-specific (*x*-axis). This pattern is not present in the latent embedding features. We hypothesize that this pattern occurs due to KPMS automatically assigning a majority of syllables to DO-specific motifs due to the larger variation in behavior of DO mice. In addition to these strain-specific behavioral features, we identified a set of shared features (distributed along the diagonal). These results suggested that there are both shared behavioral patterns associated with aging and frailty across B6 and DO mice, as well as strain-specific patterns.

These results motivated us to build separate predictive models for B6, DO, and combined B6-DO datasets in the next section to predict chronological age and frailty using supervised, unsupervised, and combined feature sets.

### 3.3 Building predictive models to predict chronological age, frailty, and proportion of life lived

We performed a series of predictive modeling experiments targeting chronological age, frailty, and the proportion of life lived (PLL; only for DO mice) to quantify the predictive utility of supervised and unsupervised behavior features. Using three distinct datasets (B6J, DO, and a combined B6J-DO cohort), we benchmarked the performance of three feature configurations: supervised (S), unsupervised (U), and a combined set (S+U).

We evaluated three predictive algorithms: a parametric linear model with an elastic net penalty to address multicollinearity, and two non-parametric tree-based ensembles - Random Forests and Gradient Boosting-to capture potential non-linearities and interactions between features. We used a repeated nested cross-validation procedure (10 × 5 with 5 repeats) to evaluate the model performance. We used the outer folds to evaluate the model performance and performed hyperparameter tuning using grid search in the inner loop. We quantified the model performance using median absolute error (MAE), root mean squared error (RMSE), and R-squared (*R*^2^). To remain consistent with our previous work, we labeled the models trained with supervised and unsupervised features as vFRIGHT, vFI, vPLL, uvFRIGHT, uvFI, and uvPLL, respectively (Figure 1B). We labeled the models trained using both feature sets (S+U) as bvFRIGHT, bvFI, and bvPLL.

Through these modeling experiments, we addressed three primary research questions:

1. *Feature sets comparison:* How does the performance of unsupervised features compare to supervised features, and does the combination of two provide an improvement in predictive accuracy?
2. *Cross-strain generalization:* Do predictive models trained on B6J using unsupervised features generalize to DO mice, and vice versa?
3. *Feature Importance and Interpretation:* Which specific set of unsupervised behavioral features is most predictive of aging and frailty across different genetic backgrounds? Which unsupervised features, associated with age and frailty, when included with supervised features, are important for predicting age and frailty?

#### 3.3.1 Feature sets comparison

To evaluate the comparative predictive utility of the feature sets, we first compared the performance of supervised (vFRIGHT) with unsupervised (uvFRIGHT) models for chronological age prediction. Overall, we found that uvFRIGHT did not outperform vFRIGHT in predicting chronological age across all three datasets (Figure 3A, D). Next, we found that a model trained on both features outperformed vFRIGHT and uvFRIGHT models in predicting chronological age across all three datasets. The combined model (bvFRIGHT) had MAEs of 10.6, 12.4, and 11.7 weeks for the B6J, DO, and B6J-DO data, respectively. This suggested that unsupervised features may identify subtle age-related behaviors not captured by supervised features alone, thereby helping to better predict chronological aging.

**Figure 3:**
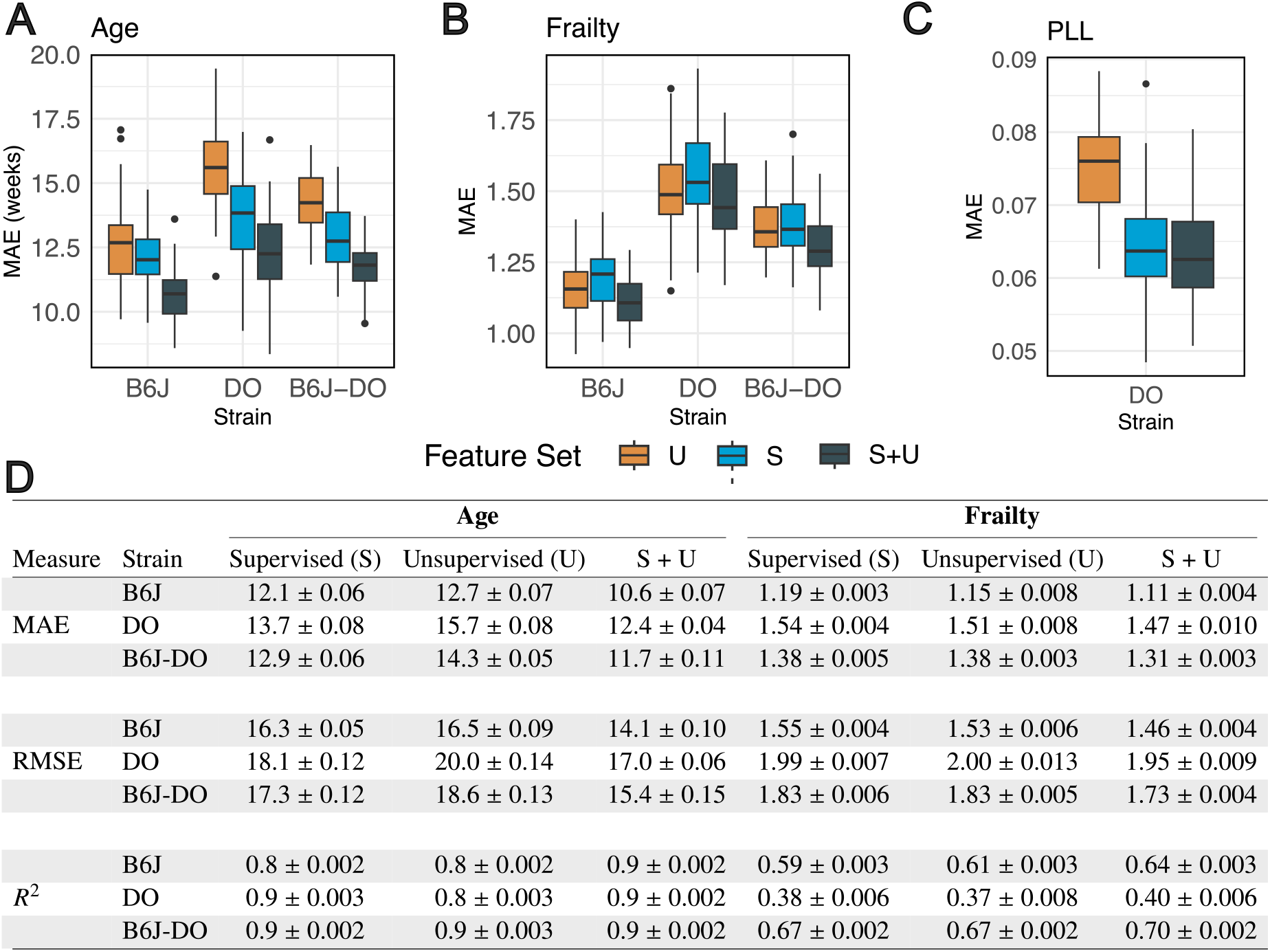
Performance of predictive models using supervised, unsupervised, and combined feature sets in predicting (A) chronological age (Age), (B) frailty, and (C) proportion of life lived (PLL) across three datasets - B6J, DO, and combined. For each dataset (x-axis) and feature set (color), we plot the performance of the best-performing model (penalized linear regression, random forest, or gradient boosting) based on median absolute error (MAE) across 50 cross-validation folds. (D) The table below shows the performance of all models across three metrics - MAE, root mean squared error (RMSE), and R-squared (*R*^2^). Lower MAE and RMSE and higher *R*^2^ indicate better model performance.

In contrast to chronological age clocks, we found that unsupervised features (uvFI) outperformed supervised features (vFI) in predicting frailty across all three datasets. Furthermore, the combined model (bvFI) outperformed vFI and uvFI across all three datasets (Figure 3B). The combined model (bvFI) had MAEs of 1.11, 1.47, and 1.31 for the B6J, DO, and B6J-DO datasets, respectively (Figure 3D). Intuitively, an MAE of 1.5 relative to manual FI items is approximately equivalent to a mis-scoring of one item by 1.0 and another by 0.5, or three items by 0.5. The improved performance of bvFI over uVFI suggested that adding expert-selected supervised behavioral features, such as gait, posture, morphometrics, and traditional open-field measures, may capture interpretable aging biomarkers missed by unsupervised algorithms.

Finally, we evaluated the performance of supervised (vPLL) and unsupervised (uvPLL) models in predicting mortality on the DO dataset. We rescaled chronological age to the proportion of life lived (PLL = Age at test/Lifespan) to remove survivorship bias across diet groups. Similar to the FRIGHT models above, we found that vPLL (MAE = 0.06, *R*^2^ = 0.33) outperformed uvPLL (MAE = 0.08, *R*^2^ = 0.17) in predicting PLL (Figure 3C). As with chronological age prediction, the combined model (bvPLL) outperformed uvPLL and vPLL, with MAE = 0.06 and *R*^2^ = 0.37, suggesting that the unsupervised features may identify behaviors related to mortality that are not captured by supervised features alone, thereby improving mortality prediction.

#### 3.3.2 Cross-strain generalization

A significant hurdle in behavioral phenotyping is the lack of generalizability of predictive models across genetic backgrounds. We previously demonstrated that supervised features (vFRIGHT) exhibit poor cross-strain generalization for chronological age (*B*6*J* → *DO*: MAE = 77.3 weeks; *DO* → *B*6*J*: MAE = 64 weeks) and frailty (vFI) (*B*6*J* → *DO*: MAE = 2.58; *DO* → *B*6*J*: MAE = 1.95). Therefore, the models trained on isogenic B6J mice failed to capture the broader behavioral phenotypic variance in DO mice, and vice versa.

To address the second question, we evaluated whether the unsupervised features could mitigate this translation gap. We found that unsupervised features also exhibited poor cross-strain generalization in predicting both chronological age (*B*6*J* → *DO*: MAE = 46.1 weeks; *DO* → *B*6*J*: MAE = 34.9 weeks) and frailty (*B*6*J* → *DO*: MAE = 2.2; *DO* → *B*6*J*: MAE = 2.4). While unsupervised models showed a marginal improvement in age-prediction error compared to supervised models, the errors remained prohibitively high for practical application across strains.

#### 3.3.3 Feature Importance and Interpretation

While our combined models (bvFRIGHT, bvFI, and bvPLL) achieved the highest predictive performance, we sought to bridge the gap between their accuracy and biological interpretability. We were faced with two primary challenges. First, the best-performing algorithms - XGBoost and Random Forest - are non-parametric ‘black boxes’ that lack the direct interpretability and transparency of coefficients found in penalized linear models^28^. Second, while unsupervised features provide high-density behavioral annotations, they are organized at a sub-second timescale^29^ and lack the inherent interpretability of supervised features. This creates a gap between predictive power and biological insight.

To address the first challenge, we took two approaches to probe the ‘black box’. First, we evaluated the accuracy of individual unsupervised feature sets (syllable usage, transitions, and latent embeddings) in isolation and compared their marginal performance with supervised features (Figure 4 A, B, and C; Table 4G). For chronological age prediction (Figure 4A), we found that, among unsupervised features, syllable usage set features were closest in performance to supervised features across all three datasets. Further, syllable transitions had greater predictive power in the DO dataset than in the B6J and B6J-DO datasets. We found a similar trend for frailty and PLL predictions: the syllable usage feature set continued to dominate the latent embeddings and syllable transitions in terms of predictive accuracy, and its performance was closest to that of supervised features. As before, the syllable transitions performed better in the DO dataset compared to the B6J and B6J-DO datasets. This approach enabled us to isolate the marginal performance of individual unsupervised feature sets for comparison with that of supervised features.

**Figure 4:**
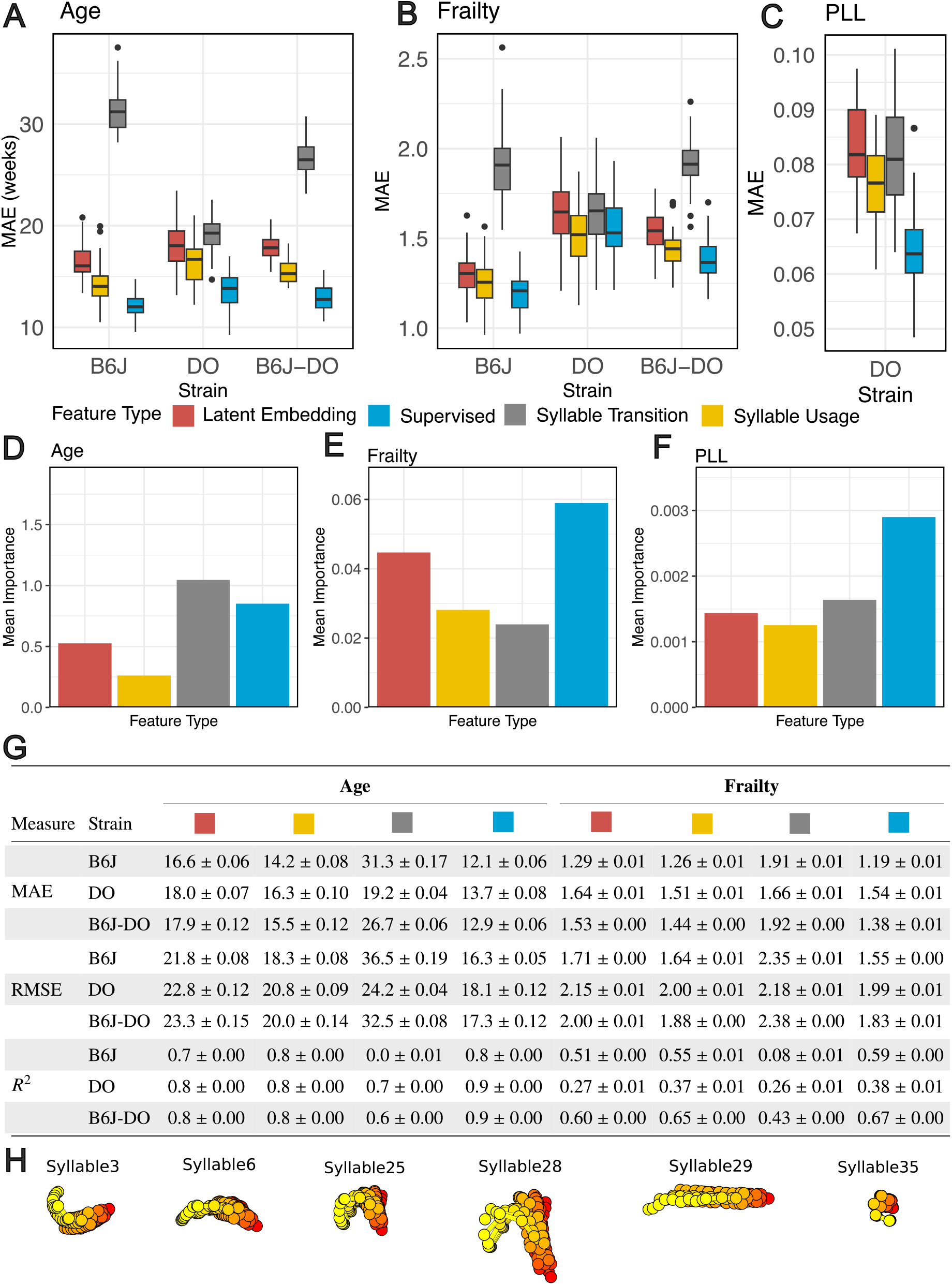
Feature importance and interpretability of supervised and unsupervised behavioral features. (A–C) Panels compare supervised features with each unsupervised category (syllable usage, syllable transitions, latent embeddings) for predicting chronological age (A), frailty (B), and proportion of life lived (C) across B6J, DO, and B6J–DO datasets; performance is reported as mean absolute error (MAE). (D–F) Permutation-based importance quantifies each feature set’s contribution to model performance for chronological age (D), frailty (E), and proportion of life lived (PLL) (F); larger values indicate a greater increase in prediction error when that set is permuted. (G) Table summarizes MAE, root mean squared error (RMSE), and *R*^2^ for unsupervised categories versus supervised features; lower MAE and RMSE and higher *R*^2^ indicate better performance. (H) Mean pose trajectories for selected high-importance KPMS syllables (see also Supplemental Movies).

Because modeling feature sets separately doesn’t account for relative contributions from different sets in the integrated (bvFRIGHT, bvFI, and bvPLL) models, we employed permutation-based feature importance in our second approach (see Methods). This allowed us to delineate the relative contributions of each feature set in the integrated (bvFRIGHT, bvFI, bvPLL) models for the B6J-DO dataset. We found that, for bvFRIGHT (Figure 4D) and bvPLL (Figure 4F), features from the syllable transition and supervised feature sets yielded the largest increase in MAE (i.e., higher mean importance) when permuted, indicating they were the most critical drivers of models accuracy. In contrast, for the biological age/frailty model (bvFI; Figure 4E), we found that the supervised feature set had the highest mean importance, followed by latent embedding, syllable usage, and syllable transition. Overall, using the first approach, we found that across predictive models for age, frailty, and PLL, supervised features had the best marginal performance, followed by the syllable-usage feature set, which had the best marginal performance among the unsupervised feature sets. Our permutation-based approach allowed us to estimate the relative contributions of the integrated (supervised + unsupervised) features in the combined dataset, revealing that supervised features dominated mean importance in vFI models and remained competitive in vFRIGHT and vPLL models.

Although the approaches above provided some model-based intuition into how feature sets contributed, they did not resolve the second challenge: insight into the behaviors underlying the unsupervised motifs. To decode unsupervised behaviors, we identified the top-ranking syllables in the syllable usage feature set via permutation-based ranking and generated representative video clips (Figure 4H; also see supplemental movie). We found that the top-ranking syllables corresponded to standard locomotory behaviors. Syllables 3, 6, 25, and 28 represent turning left and turning right at different speeds. Syllable 29 corresponds to the darting behavior, whereas syllable 35 corresponds to the animal being stationary. Notably, although turn-based behavioral features were not included in our previously published work^26, 27^, we have since developed dedicated supervised classifiers for these movements in concurrent research^30^. Our findings here demonstrate that these behaviors can be captured with high predictive utility either directly via unsupervised motifs or through dedicated supervised classifiers. Moving forward, the ability to leverage both approaches in tandem provides a robust framework for effectively bridging the gap between automated, data-driven discovery and expert-defined ethology in geroscience research.

## 4 Discussion

The manual frailty index is among the strongest predictors of morbidity and mortality in preclinical aging research, yet its reliance on subjective, expert-based scoring limits both scalability and cross-laboratory reproducibility^7, 8, 9^. In our previous studies, tester effects accounted for 42% and 18% of the variance in manual FI scores in the B6J and DO datasets, respectively; in the DO cohort, this technical variation exceeded the biological variation introduced by severe dietary restriction (14%)^14, 15^. To mitigate these limitations, we developed the visual frailty index (vFI), an objective, machine-vision-based framework that predicts frailty from supervised behavioral features extracted from open-field video^14, 15^. The vFI eliminates scorer subjectivity and provides a scalable alternative for both isogenic (B6J) and genetically diverse (DO) mouse populations.

However, a fundamental constraint of the supervised vFI is its dependence on expert-defined features (morphometric measures, gait parameters, and traditional open-field metrics), which, by design, capture only the behavioral repertoire that human observers have already associated with aging. Supervised behavior classifiers annotate only a fraction of the total video, leaving the majority of potentially informative behavioral data unanalyzed. In the present study, we asked whether unsupervised behavioral discovery via Keypoint-MoSeq (KPMS) could complement the vFI by mining this untapped behavioral space, and we organized our investigation around three questions: (1) how unsupervised features compare with supervised features for predicting aging outcomes; (2) whether unsupervised features improve cross-strain generalization; and (3) which unsupervised feature categories are most informative, and what biological behaviors they represent. We discuss each in turn.

### 4.1 Supervised and unsupervised features provide complementary predictive information

Our central finding is that combining supervised and unsupervised feature sets consistently improved prediction of chronological age, frailty, and proportion of life lived across all datasets (B6J, DO, and combined B6J-DO). The combined models (bvFRIGHT, bvFI, bvPLL) outperformed both supervised-only and unsupervised-only models, with the combined frailty model (bvFI) achieving MAEs of 1.11, 1.47, and 1.31 CFI units for the B6J, DO, and B6J-DO datasets, respectively. In practical terms, an MAE of approximately 1 CFI unit is equivalent to a single FI item being mis-scored by one point, or two items being mis-scored by 0.5 points, a level of error comparable to or smaller than the inter-rater variability observed in manual FI assessment^8, 14^.

The consistent improvement from feature fusion, though modest in absolute terms, held across all three prediction targets and all three datasets, lending confidence that the two feature sets capture partially non-overlapping information about aging-related behavioral change. Supervised features (gait speed, stride regularity, body morphometrics, and spinal flexibility) provide biologically grounded, interpretable metrics with well-established links to musculoskeletal and neuromuscular decline^14, 31^. Unsupervised KPMS features, by contrast, summarize the full temporal structure of the animal’s behavioral repertoire without prior assumptions, and may capture subtle postural or kinematic patterns, such as altered behavioral sequencing or shifts in the time allocation across motor programs, that do not map neatly onto any single expert-defined behavior.

For chronological age and PLL, supervised features outperformed unsupervised features. For frailty, unsupervised features performed comparably to supervised features, with a small numerical advantage (e.g., B6J MAE: S = 1.19 vs. U = 1.15; a difference of ∼0.04 CFI units). While this pattern was consistent across all three datasets, the magnitude of the difference is small relative to the measurement scale, and the two feature sets should be considered broadly equivalent for frailty prediction. The key result is not that one approach dominates the other, but rather that their combination yields consistent improvements, indicating that the two feature sets capture partially non-redundant aspects of aging-related behavioral change.

An intriguing pattern in our results is that unsupervised features performed relatively better for frailty prediction than for chronological age prediction, where supervised features held a clearer advantage. One possible explanation is that chronological age tracks a monotonic trajectory well captured by progressive, easily quantifiable declines in locomotion and body composition, precisely the features targeted by the supervised set. Frailty, by contrast, reflects a multisystem vulnerability that may manifest through subtler reorganizations of the behavioral repertoire (changes in the temporal structure, sequencing, or relative allocation of motor programs) that are better captured by the holistic, data-driven segmentation of KPMS. This hypothesis is consistent with the clinical frailty literature, in which frailty is understood as a state of reduced physiological reserve that affects multiple systems simultaneously and need not be captured by any single physical measure^1, 3, 5^.

### 4.2 Cross-strain generalization failure reveals population-specific behavioral aging signatures

Neither supervised nor unsupervised features resolved the challenge of cross-strain generalization. Models trained on B6J performed poorly when applied to DO mice, and vice versa, regardless of feature set. This failure is consistent with our previous finding that population-specific models are required for accurate frailty prediction^15^ and extends it by demonstrating that the generalization barrier is not an artifact of the supervised feature selection: it persists even when the feature space is learned entirely from data.

This result has important biological implications. The behavioral manifestations of aging are not universal across genetic backgrounds; rather, they are shaped by the interaction between genotype and the aging process. Isogenic B6J mice age through a relatively constrained behavioral trajectory, whereas DO mice, with their high genetic diversity, exhibit a much broader landscape of aging-associated behavioral phenotypes. KPMS automatically assigned a disproportionate number of syllables to DO-specific motifs (Figure 2D,E), reflecting the greater behavioral heterogeneity of the outbred population. These DO-specific syllables were often among the most predictive features for DO models, but carried no predictive value for B6J mice.

From a practical standpoint, this finding reinforces the principle that video-based aging clocks, like other biomarkers, must be calibrated to the target population. We expect that expanding the training data to include additional genetic backgrounds—such as UM-HET3 mice used by the Interventional Testing Program (ITP)^32, 33^ or BXD recombinant inbred strains^34^—will be necessary for building broadly applicable tools. However, as we noted previously^15^, there is likely a precision–generality tradeoff: models trained on more diverse cohorts may gain accuracy across populations at the cost of precision within any single strain. Characterizing this tradeoff systematically is an important direction for future work.

### 4.3 Syllable usage features are the most predictive unsupervised category, and top syllables correspond to canonical behaviors

Among the three categories of unsupervised features (syllable usage, syllable transitions, and latent embeddings), syllable usage was consistently the most predictive for age, frailty, and PLL across all datasets (Figure 4A–C, G). This finding suggests that aging-related behavioral change is primarily reflected in *how much time* an animal spends in different motor programs, rather than in the *sequential structure* of transitions between them or in the continuous latent pose space. This is biologically intuitive: an older, frailer mouse is expected to spend more time in stationary postures and less time in vigorous locomotion, a shift that is directly captured by syllable usage proportions.

Permutation-based feature importance analysis of the combined models revealed that supervised features retained the highest average importance, with syllable usage and syllable transition features providing complementary contributions (Figure 4D–F). This pattern suggests that the combined models leverage supervised features as the primary predictive scaffold, supplemented by targeted unsupervised features that capture residual variance not explained by the expert-selected set.

To gain biological insight into the unsupervised features, we examined the top-ranking KPMS syllables via their average pose trajectories and representative video clips (Figure 4H; Supplemental Movies). The most important syllables corresponded to recognizable motor patterns: turning at various angles and speeds (syllables 3, 6, 25, 28), straight forward locomotion (syllable 29), and a stationary resting pose (syllable 35). These behaviors overlap with the supervised feature categories (angular velocity, speed, rearing/inactivity), suggesting that the predictive contribution of unsupervised features arises less from the discovery of entirely novel behavioral categories and more from capturing finer-grained kinematic variation within canonical behaviors—for example, distinguishing slow from fast turns, or quantifying precise time allocation across motor states at a temporal resolution that the supervised feature set does not explicitly encode. In this view, the value of unsupervised segmentation lies not in replacing expert-defined features but in providing a complementary, higher-resolution decomposition of the same underlying behavioral space.

### 4.4 Practical accessibility and community resources

The ease of use of automated behavioral phenotyping tools for non-computational laboratories remains a critical barrier to adoption in geroscience^15^. To lower this barrier, we provide: (1) a KPMS model pre-trained on the combined B6J-DO population, enabling other groups to extract unsupervised features from their own open-field videos without retraining; (2) population-specific aging clocks (supervised, unsupervised, and combined) for B6J, DO, and combined datasets; and (3) we release the entire (1138 tests) open field video and manual frailty dataset for future analysis and algorithm development. For supervised feature extraction, we use the JAX Animal Behavior System (JABS)^30^, which provides an integrated pipeline for tracking, pose estimation, feature extraction, and behavior classification.

### 4.5 Limitations

Several limitations of this study should be acknowledged. First, the KPMS training set was limited to 150 videos due to computational memory constraints rather than a principled analysis of sufficiency. While the stratified sampling procedure ensured balanced representation across ages, sexes, strains, and diets, we did not perform a learning-curve analysis to verify that 150 videos is sufficient for stable downstream prediction performance. There is a sex imbalance in the DO cohort (predominantly female) due to limitations of the DO population^15^, which precluded a systematic analysis of sex-specific aging trajectories, which is an important dimension given the known sex differences in frailty and lifespan^14^.

### 4.6 Future directions

An important extension of the present work is to lower the barrier of digital phenotyping for the biomedical community. We recently introduced a home cage monitoring system for the laboratory mouse that enables continuous 24/7 observation of group-housed animals^35^. In addition, the field has several other platforms for continuous monitoring. Building a vFI for the laboratory mouse in these form factors would greatly lower the barrier of these advanced methods. Video-based home-cage monitoring is a rapidly advancing field^36, 37, 38^ that could, in principle, provide continuous, longitudinal frailty assessment throughout an animal’s life without the need for repeated handling^39, 40^. Data from social interactions, circadian activity patterns, sleep-wake states, feeding and drinking behavior, and nesting, combined with the gait, postural, and morphometric features described here, could enable a richer and more ecologically valid assessment of biological aging. Unsupervised behavioral segmentation methods like KPMS are particularly well-suited to this setting, where the behavioral repertoire is broader and less constrained than in the open field, and where human-defined behavioral categories are even less likely to be comprehensive.

More broadly, the complementarity of supervised and unsupervised features demonstrated here motivates a general framework for behavioral biomarker discovery in aging research: begin with expert-defined, biologically grounded features to establish an interpretable baseline, then augment with data-driven features to capture residual behavioral information that escapes human categorization. As pose estimation and behavioral segmentation tools continue to mature, and as standardized video phenotyping platforms become more widely available, this integrated approach has the potential to transform aging research from labor-intensive, subjective assessments to scalable, objective, and continuously updated measures of biological age.

## 5 Methods

### 5.1 Open field testing

All animals and methods have been previously described. We carried out behavior tests according to published methods^16, 17, 41, 42^. Our open field arena measures 52 cm by 52 cm by 23 cm. The floor is white PVC plastic and the walls are gray PVC plastic. To aid in cleaning maintenance, a 2.54 cm white chamfer was added to all the inner edges. Illumination is provided by an LED ring light (Model: F&V R300). The ring light was calibrated to produce 400 − 600 lux of light in each of our 24 arenas. Tests were performed between ZT10 and 16.

### 5.2 Animals

Both B6J and DO datasets have been previously described^26, 27, 43, 44^. The imbalance in the male/female ratio in this study is by design. We observed high rates of fighting in DO males, particularly as they aged. This made long-term aging studies in DO males challenging. Therefore, early aging studies in DO almost exclusively used females, including^44^ whose animals are repurposed for this paper. We have since solved the aggression problem in DO males through appropriate enrichment. In this study, we do not analyze differences in males and females due to this imbalance and lack of power.

### 5.3 Frailty Index Items

The JAX Nathan Shock Center FI procedures have evolved over time. The initial data used in^26^ used 27 ordinal items (0,0.5,1 scale) and two continuous measures (temperature and weight). Continuous measures are collected but not used in the final manual FI score. More recently, the JAX NSC has expanded the number of items to 30 ordinal items in a fragility index^43, 44^. Since we wished to compare the B6J and DO clocks fairly, we only used the 27 common FI items. The full frailty items for DO mice are reported in^43^ and^44^.

### 5.4 Behavior Segmentation and Feature Extraction

For each of the three populations of mice (B6J only, DO only, and B6J and DO combined), we trained a Keypoint-MoSeq (KPMS) model to extract behavioral syllables present in the video data. Each syllable roughly corresponded to a distinct mouse action, such as walking, grooming, or rearing. Keypoint-MoSeq uses a Switching Linear Dynamical System to model behavior and produces, for each frame of the input video, a representation in a lower-dimensional pose space as well as a discrete syllable assignment. Importantly, the number of latent dimensions and syllables is learned automatically from the data.

To ensure a balanced representation of both chronological and biological ages in the training data, we used stratified sampling to select 150 videos for training the combined model. This number of videos was chosen because it was the maximum number of videos that could simultaneously fit in memory during the KPMS training process. Empirically, we found that this value depended on both the size of each video as well as the latent dimension computed during the data embedding preprocessing step. The sampled videos were selected to encapsulate the distribution of mice across all ages, sexes, diets, and strains (Figure S1A). Stickiness parameters of κ = 10^6^ and κ_red_ = 10^5^ were chosen for the initial Autoregressive-Hidden Markov Model fitting and full model fitting, respectively, to ensure that the resulting syllables had a median length of approximately 0.5 s or 15 frames. For each dataset, we trained 6 KPMoSeq models with different starting seeds and selected the model with the highest Expected Marginal Likelihood (EML) score for downstream analysis. We direct readers interested in the KPMS training process to the official documentation.

Using both the continuous low-dimensional pose representation and the discrete syllable assignments from the KPMS behavior segmentation, we extracted three classes of features: the mean, median, and standard deviation along each dimension of the latent pose representation; the proportion of time spent in each syllable; and the desparsified transition probabilities between syllables.

### 5.5 Metasyllable Construction

Due to the large number of syllables, we found the transition matrix features to be noisy and sparse, particularly between rare syllables (Figure S1C). To reduce noise and increase the information gained from transition probabilities, we clustered similar syllables and grouped them into 5 “meta-syllable”. Meta-syllables were created by building a dendrogram by iteratively merging pairs of syllables with largest cosine similarity between median syllable trajectories. The dendrogram was then broken at a *y*-level to create 5 components corresponding to the metasyllables. Each transition between syllables was grouped into the corresponding metasyllable transitions, which were used as features for downstream regression. In practice, we found that including the full, sparse transition matrix in the feature set did not significantly affect the performance of the downstream prediction tasks.

### 5.6 Predictive Modeling

We estimated tester effects by modeling the cumulative frailty index using a linear mixed model with tester and batch as random effects, and age (in weeks), weight, and sex as fixed effects. We used restricted maximum likelihood (REML) to estimate variance components and best linear unbiased predictors (BLUPs) to estimate tester- and batch-specific effects. We subtracted the estimated tester and batch effects from the cumulative FI scores. We labeled them as the tester- and batch-adjusted FI scores, 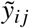.

For the cross-validation predictive modeling experiments, we ensured that repeat measurements from the same mouse were assigned to either the training or test fold, but not both.

To assess the relative contribution of each feature to our non-parametric models, we utilized permutation-based feature importance^45^. This method measures the marginal increase in prediction error (MAE) following the stochastic shuffling of a features values. By permuting the feature, the inherent relationship between the predictor and the target - either chronological age or frailty or proportion of life lived - is effectively severed. A significant increase in the resulting error indicates that the model relies heavily on that feature for accuracy, whereas a negligible change suggests the feature provides redun-dant or minimal predictive information. For each feature *f* across fold *k*, we computed its importance score as the difference between the model’s mean absolute error (MAE) on the *k*^th^ test fold and the MAE after permuting feature *f*, MAE(*f*_permuted_):

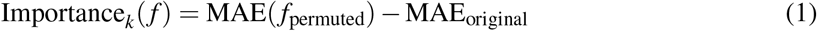

We averaged the importance scores across all folds to obtain a final importance score for each feature:

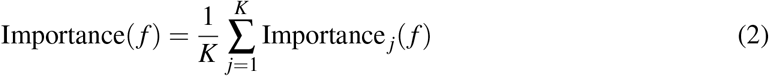

where *K* is the total number of folds in the cross-validation. We averaged the importance scores for features in the same category (e.g., syllable usage, transition, latent embeddings, supervised) to otain the overall importance of each feature category.

## 6 Acknowledgments

We thank Gary Churchill for critical feedback on the manuscript. We thank Camille Berger-Liedtka for project management and support. We thank Kumar Lab members for feedback and helpful discussions.

## 7 Author contributions

Data were from previously published studies^14, 15^. G.S.S. and V.K. conceived the project. D.M.M. and G.S.S. implemented the software and contributed to different components of the model training and experimental analyses. G.S.S. and V.K. supervised the project. All authors contributed to writing, editing, and reviewing the manuscript. V.K. secured funding for the project.

## 10 Data availability

All data will be made available at ADDURLHERE.

## 11 Code availability

All source code for training B6J-DO Keypoint-MoSeq model and agin-clock models is available at https://github.com/KumarLabJax/B6JDO-KPMS-Pipeline and https://github.com/KumarLabJax/Unsupervised-vFI respectively.

## 12 Supplement

**Table S1:**
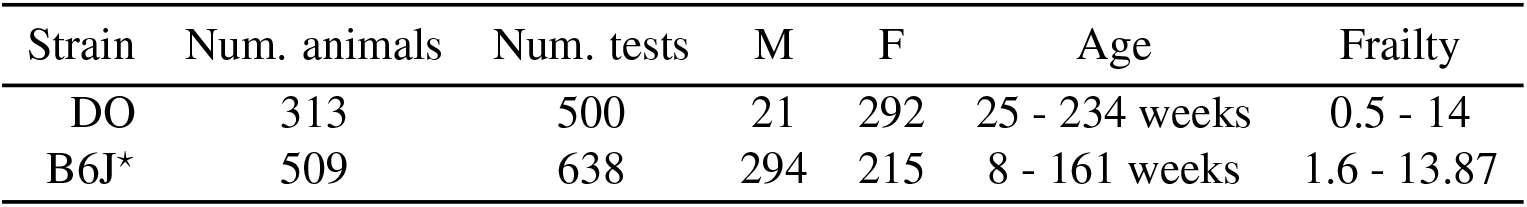
Summary table of animals and frailty scores in our dataset. ⋆The B6J dataset differs slightly from that used in^26^ owing to minor updates to the dataset. This difference does not affect the overall conclusions.

| Strain | Num. animals | Num. tests | M | F | Age | Frailty |
| --- | --- | --- | --- | --- | --- | --- |
| DO | 313 | 500 | 21 | 292 | 25 - 234 weeks | 0.5 - 14 |
| B6J* | 509 | 638 | 294 | 215 | 8 - 161 weeks | 1.6 - 13.87 |

**Figure S1:**
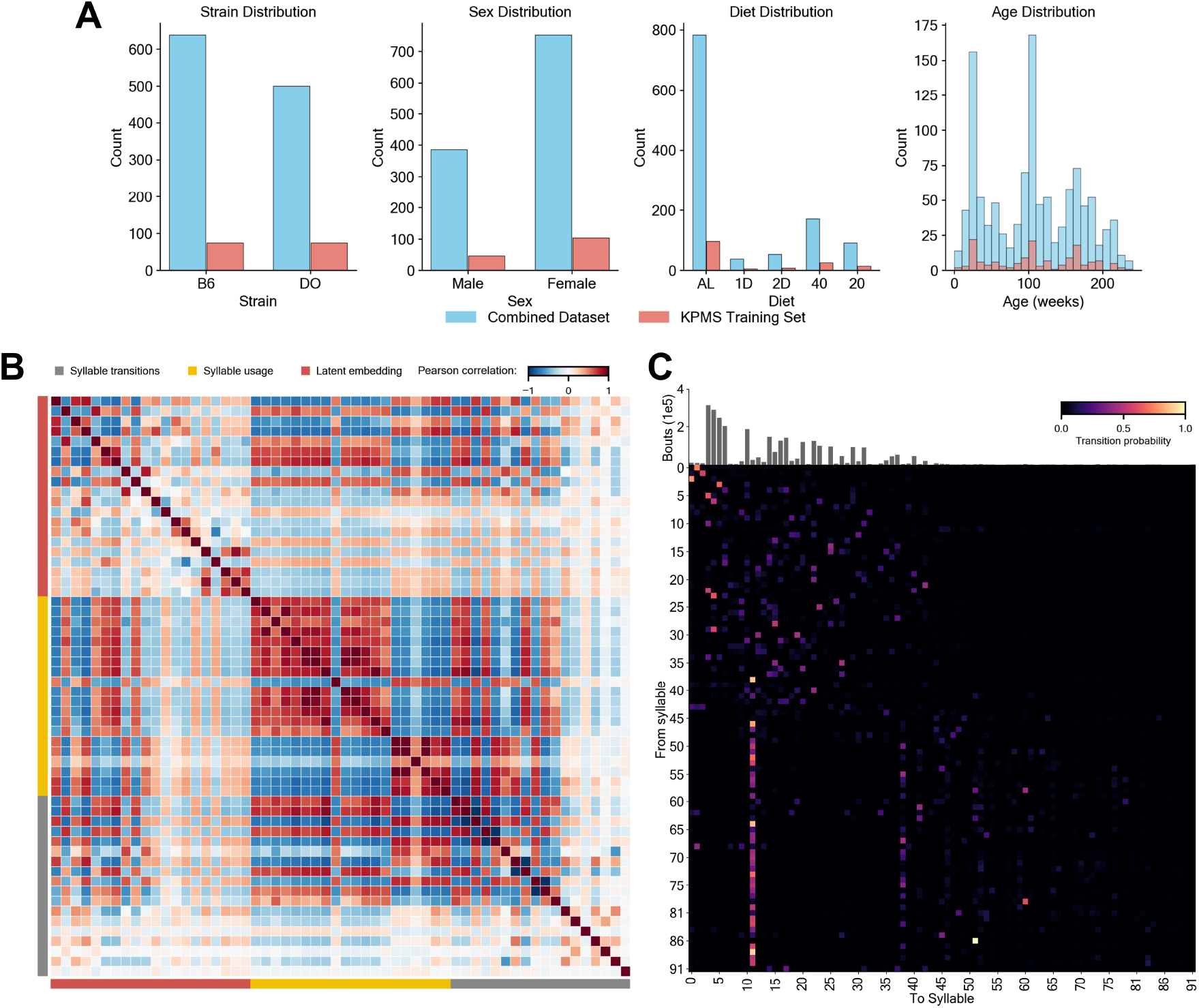
Keypoint-MoSeq Training and Inference. (A) Distribution of Combined B6 + DO dataset and KPMS training set (see Table 1) for strain, sex, diet, and age. (B) The estimated Pearson correlation matrix across all video features captures the linear interdependence (relationship) across multiple video features in our dataset. The the off-diagonal elements capture the correlation between each pair (row, column) of color-coded variables. Only the 20 features from each unsupervised aging feature type with highest magnitude of Pearson correlation with frailty are depicted. (C) The sparse mean transition matrix between all syllable pairs. Each row was normalized to sum to one to resemble a probability distribution. The histogram on the top displays the total number of bouts across the entire dataset per feature.

